# Model-free decision making resists improved instructions and is enhanced by stimulus-response associations

**DOI:** 10.1101/2022.11.23.517672

**Authors:** Raúl Luna, Miguel A. Vadillo, David Luque

**Author notes:** Corresponding authors. E-mail address (Raúl Luna) E-mail address (David Luque).

## Abstract

Human behaviour may be thought of as supported by two different computational-learning mechanisms, model-free and model-based respectively. In model-free strategies, stimulus-response associations are strengthened when actions are followed by a reward and weakened otherwise. In model-based learning, previous to selecting an action, the current values of the different possible actions are computed based on a detailed model of the environment. Previous research with the two-stage task suggests that participants’ behaviour usually shows a mixture of both strategies. But, interestingly, a recent study by da Silva and Hare (2020) found that participants primarily deploy model-based behaviour when they are given detailed instructions about the structure of the task. In the present study, we reproduce this essential experiment. Our results confirm that improved instructions give rise to a stronger model-based component. Crucially, we also found a significant effect of reward that became stronger under conditions that favoured the development of strong stimulus-response associations. This suggests that the effect of reward, often taken as indicator of a model-free component, is related to stimulus-response learning.

## 1. Introduction

It is often assumed that behaviour is based on two types of processes: goal-directed and habitual. From a computational point of view, each of these processes has been related to two different reinforcement-learning (RL) strategies: model-free and model-based, respectively (Daw, Niv & Dayan, 2005). Following these authors, in the case of model-free strategies, stimulus-response (S-R) associations are strengthened when actions or responses are followed by a reward and become weakened otherwise (Sutton & Barto, 1988). On the other hand, model-based learning generates behaviour determined by the ongoing values of all available actions. The computation of these values is based on a model of the environment, that is to say, a sort of “cognitive map” in a non-spatial domain (Dolan & Dayan, 2013). In contrast to model-free representations, these maps consider not only if current actions lead to immediate rewards, but also if they lead to new states in which other actions may produce other (better) rewards.

There is an extensive literature exploring the model-free vs. model-based dichotomy using a particular experimental paradigm: the two-stage task or two-choice Markov decision task (Daw, Gershman, Seymour, Dayan & Dolan, 2011; Decker, Otto, Daw & Hartley, 2016; Miller, Botvinick & Brody, 2017; Kool, Gershman & Cushman, 2018). Each trial in this task requires participants to go through two sequential stages. In the first stage participants are asked to select one of two options. This choice is followed by a second stage with two possible scenarios or states. In most trials, a specific option in the first stage causes a transition to a determined state in the second stage (i.e., a common transition), but in a minority of trials it may also cause a transition to the alternative state (i.e., a rare transition). At the second stage, participants are asked again to choose between two options, each leading to a different reward. The specific options presented to participants during the second stage depend on the scenario or state in which the second stage takes place. Model-free strategies lead to the repetition of actions that have previously been rewarded, regardless of the type of transition that brought the participant to a certain second-stage state in past trials. Because model-based strategies include knowledge about whether rewards were obtained as a result of an unlikely transition, they can lead to the selection of the opposed first-stage action in future trials to obtain the same reward in the second stage. Therefore, actions not leading to reward in a current trial may still be executed in future trials if the transition was rare.

Because model-free agents are prone to repeating a first-stage action that ended up in a reward irrespective of the transition they experienced, these are expected to show a positive main effect of reward in the previous trial. In contrast, model-based agents are expected to show a Reward X Transition interaction. This is because, based on a cognitive map of the task, the most rational decision is to select the first-stage action that will most likely lead to the largest second-stage reward. Of course, it is possible to combine both strategies. Such “hybrid” agents should show both a main effect of reward and a Reward X Transition interaction. Past studies have revealed the widespread use of hybrid strategies in healthy adult humans (Daw et al., 2011; Decker et al., 2016). More specifically, research suggests a prevailing use of model-based strategies, but also a significant presence of model-free behaviour, even under experimental conditions specifically designed to favour model-based learning (Kool, Cushman & Gersham, 2016; Kool, Gershman & Cushman, 2017; Kool, Gershman & Cushman, 2018).

Contrary to these findings, da Silva & Hare (2020) (see also da Silva, Lombardi, Edelson & Hare, 2023) demonstrated that it is possible to observe performance consistent with the use of primarily model-based strategies when participants are provided with accurate instructions about the task, so that they can create a correct cognitive model about it. That is, these authors argue that the model-free component evidenced in past studies is the result of an inaccurate internal model of the task, produced by a poor understanding of the instructions and the experimental paradigm. In addition to gathering empirical evidence supporting this view, they also conducted a computational-modelling analysis showing that an incorrect internal model of the task can give rise to the main effect of Reward that is often taken as evidence of model-free strategies.

In their version of the two-stage task, da Silva & Hare (2020) used first- and second-stage options that randomly swapped sides from trial to trial. This aspect of the procedure may prevent the formation of strong associations between these stimuli and specific motor commands (Molinero et al., 2021; Verleger et al., 2016, 2018). This is an aspect that may hinder the execution of habitual responses. Consistent with this, Hardwick, Forrence, Krakauer & Haith, 2019, found that training specific S-R associations always executed with the same motor command produced habits (but see Buabang et al., 2023). Also, Neal, Wood, Wu & Kurlander, 2011, showed that previously formed habits disappear when changing the response pattern. Also, Luque et al. (2020) have shown that habits formed after extended and consistent S-R training interfere with new S-R mappings. Therefore, changing response option positions at random, as it is usual in the two-stage task, may favour the operation of the goal-directed system and overshadow any possible implication of the habit system. A question that remains unsolved is whether presenting the response options at fixed locations throughout the two-stage task would enhance model-free learning—even when the participants are provided with detailed instructions so they have an accurate internal model of the task.

In the present study, we attempted to replicate da Silva & Hare’s (2020) results using the same task and the same improved instructions. Furthermore, and for the first time, we tested whether displaying response options at fixed locations leads to stronger evidence associated to model-free strategies.

In line with da Silva & Hare (2020), improved instructions did induce a larger Reward X Transition interaction effect (traditionally equated to model-based learning) than that evidenced in classical results (e.g. Daw et al., 2011; Decker et al., 2016). However, our results also show a significantly larger Reward component (traditionally equated to model-free learning) than the one in da Silva & Hare (2020). In addition, we provide evidence that fixing the location of response options potentiates the effect of Reward in participants’ behaviour, suggesting that this effect is related to S-R learning. Importantly, the methods and analysis plan of the present studies were pre-registered before data collection (https://osf.io/x9sya).

## 2. Results

As explained above, one objective of this study was to replicate the original results from da Silva & Hare (2020), that is, we sought to find evidence of increased model-based behaviour in the two-stage task with improved instructions. Additionally, this study aimed to test whether fixing the state option locations across trials potentiates an effect of reward (traditionally thought of as a model-free component) in the same task, even with detailed instructions. Two experimental groups, Replica and Fixed-Locations, were formed to achieve these objectives. The only difference between them was that, in the Fixed-Locations condition, the location of the two response options in each state remained unchanged across trials, whereas in the Replica condition, locations changed randomly.

### 2.1. Logistic regression analysis

Consecutive trial pairs were analyzed through logistic regression, where the probability of repeating the same first-stage action as in the previous trial (i.e., the probability of “staying”) was a function of reward and the type of transition in the previous trial. Reward was coded as +1 if the previous trial had been rewarded and −1 otherwise. Transition was coded as +1 if the participant’s response in stage 1 had led to the common state in stage 2 (i.e., common transition) and −1 otherwise (i.e., rare transition). Figure 1A displays the predicted stay probabilities separately for each Reward and Transition condition, both for the two groups of the present study and for the original magic carpet experiment by da Silva and Hare (2020), as well as the study by Kool et al. (2016) using the original two-stage task paradigm by Daw et al. (2011). Figure 1B shows the estimated logistic regression model coefficients for each study. According to da Silva & Hare (2020), the Reward X Transition interaction, traditionally linked to model-based behaviour, increases when participants have a good mental representation of the two-stage task induced by improved instructions. On the contrary, the main effect of reward has traditionally been thought of as evidence of a model-free strategy (Daw, Niv & Dayan, 2005).

**Figure 1.**
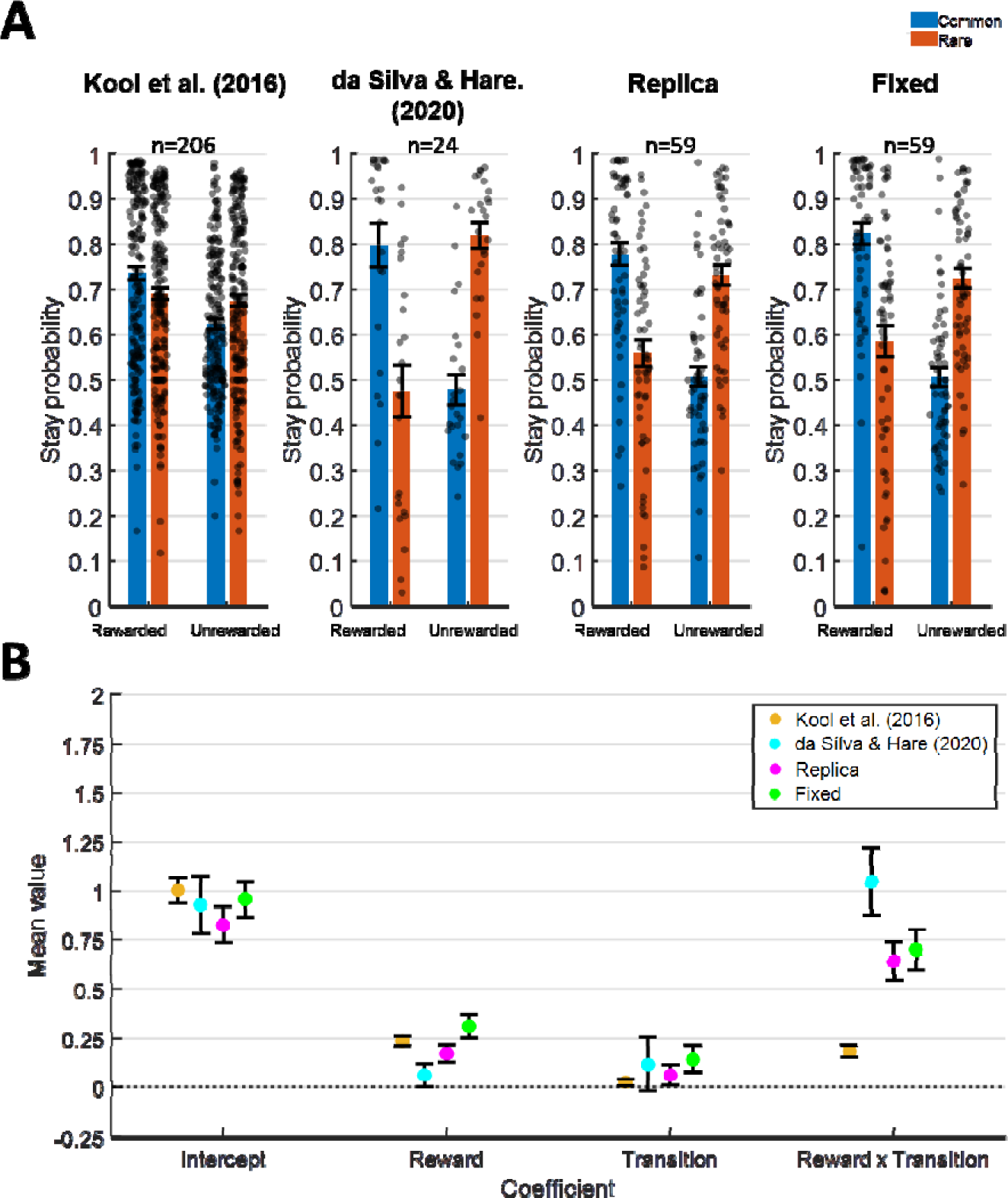
Results of the logistic regression analysis. A. Stay probabilities (probability of repeating the same response as in the previous trial) are shown in the cases when the transition in the previous trial had been Common (blue) or Rare (red). Results further distinguish whether the previous trial had been Rewarded or Unrewarded. The leftmost panel shows the results from the study by Kool et al. (2016) using the two-stage task paradigm by Daw et al. (2011); next, the results from the original magic carpet experiment in da Silva & Hare (2020) are shown; the next panel shows the results for the Replica condition and the rightmost panel shows them for the Fixed-Locations condition. Individual results are shown as well as the mean ± SEM. B. Coefficients for each of the logistic regression parameters, which were used to calculate the stay probabilities shown in the upper panels. The mean ± SEM is depicted.

As can be seen in Figure 1, the coefficient value of the Reward X Transition interaction was substantially lower in our Replica condition than in the original study by da Silva & Hare (2020), as shown by an independent samples t-test (t(81)=-2.3828, p=0.0195, two-tailed, d=-0.5553, 95% CI [-0.7819, −0.0703]). We did not find significant differences between both studies in any of the other coefficients (Intercept: t(81)=-0.7158, p=0.4761, two-tailed, d=-0.1723, 95% CI [-0.4636,0.2183]; Reward: t(81)=1.3664, p=0.1756, two-tailed, d=0.3417, 95% CI [-0.0496, 0.2671]; Transition: t(81)=-0.4145, p=0.6796, two-tailed, d=-0.0889, 95% CI [-0.2892, 0.1895]). We further tested which coefficients statistically differed from zero. This would indicate a main effect of such coefficients. Table 1 shows the results from one-sample t-tests. In the study by da Silva & Hare (2020), no main effect of Reward was observed, while there was a main effect of Reward X Transition. Our Replica study shows both significant Reward and Reward X Transition coefficients. A similar pattern can be appreciated in the study by Kool et al. (2016).

**Table 1.** Two-tailed one-sample t-tests for each of the coefficients, Intercept, Reward and Transition and Reward X Transition. Results are shown for all four studies, Kool, et al. (2016), da Silva & Hare (2020), Replica and Fixed. The values of the t-statistics and their degrees of freedom are displayed, as well as p-values and 95% confidence intervals. p-values lower than 0.05 are marked in bold, indicating those cases where a coefficient significantly differs from zero.

|  |  | Kool et al. (2016) | dS & H (2020) | Replica | Fixed |
| --- | --- | --- | --- | --- | --- |
| Intercept | t(df) | t(205)=15.0920 | t(23)=6.3203 | t(58)=8.8338 | t(58)= 10.5500 |
|  | p | **p<0.0001** | **p<0.0001** | **p<0.0001** | **p<0.0001** |
|  | 95% CI | [0.8744,1.1372] | [0.6259,1.2349] | [0.6247,0.9907] | [0.7839,1.1510] |
| Reward | t(df) | t(205)=8.9397 | t(23)=1.0656 | t(58)=3.8708 | t(58)= 5.0041 |
|  | p | **p<0.0001** | 0.2977 | **0.0002** | **p<0.0001** |
|  | 95% CI | [0.1844,0.2887] | [-0.0599,0.1871] | [0.0832,0.2615] | [0.1841,0.4296] |
| Transition | t(df) | t(205)=1.3947 | t(23)=0.8453 | t(58)=1.3282 | t(58)= 2.0374 |
|  | p | 0.1646 | 0.4067 | 0.1893 | **0.0462** |
|  | 95% CI | [-0.0105,0.0615] | [-0.1715,0.4086] | [-0.0348,0.1721] | [0.0024,0.2697] |
| Reward × Transition | t(df) | t(205)= 6.7859 | t(23)= 6.1931 | t(58)=6.8477 | t(58)= 6.6491 |
|  | p | **p<0.0001** | **p<0.0001** | **p<0.0001** | **p<0.0001** |
|  | 95% CI | [0.1321,0.2402] | [0.6988,1.3997] | [0.4410,0.8053] | [0.4848,0.9025] |

When we compared our coefficients from the Replica study with those by Kool et al. (2016), we found statistically significant differences in the Reward X Transition coefficient (t(264)=6.3894, p<0.0001, two-tailed, d=0.8004, 95% CI [0.3161, 0.5978]). That is, improved instructions boost model-based behaviour, as measured by the Reward X Transition interaction. We did not find significant differences between our Replica condition and the study by Kool et al. (2016) in the rest of the coefficients (Intercept: t(264)=-1.3093, p=0.1916, two-tailed, d=-0.2116, 95% CI [-0.4461,0.0898]; Reward: t(264)=-1.1125, p=0.2669, two-tailed, d=-0.1722, 95% CI [-0.1710, 0.0475]; Transition: t(264)=0.8415, p=0.4008, two-tailed, d=0.1110, 95% CI [-0.0500, 0.1246]).

The comparison of our Replica and Fixed-Locations studies revealed a stronger effect of Reward in the latter, as evidenced in the value of the Reward coefficient, which was larger in Fixed-Locations than in Replica (t(116)=-1.7748, p=0.0393, one-tailed, d=-0.3268, 95% CI (-∞, −0.0088]). No significant differences between the studies were found in the rest of the coefficients (Intercept: t(116)=-1.2335, p=0.2199, two-tailed, d=-0.2271, 95% CI [-0.4162, 0.0967]; Transition: t(116)=-0.7980, p=0.7868, one-tailed, d=-0.1469, 95% CI [-0.2074, ∞); Reward X Transition: t(116)=-0.5091, p=0.6942, one-tailed, d=-0.0937, 95% CI [-0.3000, ∞))^1^.

It could be argued that despite our use of improved instructions, some participants might still have failed to understand the structure of the task. Consequently, their behaviour under an inaccurate cognitive map of the two-stage task may have biased our results towards a larger effect of Reward. To address this possibility, participants were asked to complete a questionnaire at the end of the task (See the “5. Materials and Methods” section, “5.5. *Procedure*”). To rule out the possibility that our results are biased by the inclusion of participants who did not understand the instructions, in Appendix A we report additional logistic regression analyses excluding participants who failed any question in any of the questionnaires. We apply this same exclusion criterion to the participants in the experiment by da Silva & Hare (2020). The exclusion of these participants did not make a meaningful difference in the results.

### 2.2. Hybrid reinforcement learning model fits

To further analyze the extent to which participants showed model-based vs model-free behaviour in the two-stage task, we fitted the standard hybrid reinforcement learning model proposed by Daw et al. (2011) to their data. This model combines the model-free SARSA (λ) algorithm with a model-based learning algorithm, weighted by parameter *w* (0≤*w*≤1). This parameter can be interpreted as a model-based weight, with a value of 1 indicating the use of purely model-based strategies and 0 indicating a sole model-free behaviour. Figure 2 shows the estimated *w* weights for the magic carpet experiment in da Silva & Hare (2020), as well as our the Replica and Fixed-Locations studies. (See Appendix C for estimates of the rest of the parameters as well as the negative log-likelihood, whose value is minimized during model fitting). Our Replica study shows a *w* weight that is significantly lower than the one from da Silva & Hare (2020) (: t(81)=3.1706, p=0.0021, two-tailed, d=0.8209, 95% CI [0.0941, 0.4112]). The model suggests a hybrid model-free/model-based behaviour. On a different note, the hypothesis that the Fixed-Locations condition promotes the use of model-free strategies is not clearly supported by the hybrid reinforcement learning model fits. Although the model-based weight in the Fixed-locations condition was slightly lower than in the Replica condition—indicating a somewhat larger model-free behaviour—the difference between both fails to reach statistical significance (: t(116)=0.1379, p=0.4453, one-tailed, d=0.0269, 95% CI [-0.1014,)).

**Figure 2.**
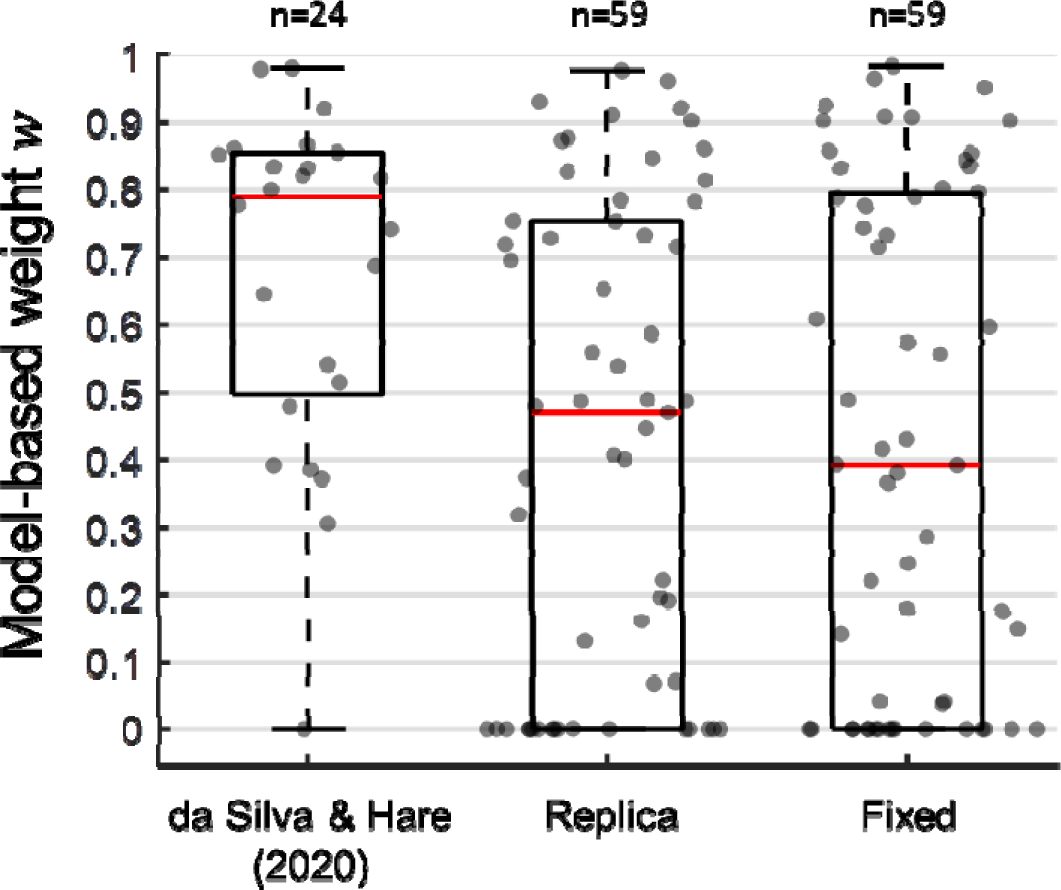
25%, 50% (median), and 75% percentiles of estimated parameters from the hybrid reinforcement learning algorithm as well as individual estimates. Data is shown for the *w* model-based weight parameter.

In sum, differences between both groups can be detected in the Reward coefficient from the logistic regression analysis, which may be compatible with differences in model-free behaviour; however, this pattern is not found in the hybrid reinforcement learning model fits (i.e., in the parameter). This may be the consequence of the single parameter not being as sensitive as the multiple logistic regression coefficients for discriminating model-free vs model-based behaviour. In line with this interpretation, da Silva & Hare (2020) concluded that the logistic regression model is better than the hybrid model at explaining first-stage choices in this task. Consequently, the reported differences between Replica and da Silva & Hare (2020) using the hybrid reinforcement learning model must be interpreted with caution.

## 3. Discussion

The present study attempted to replicate the results by da Silva & Hare (2020 Experiment 1—the experiment using the magic carpet task. See also da Silva et al., 2023), who found that, when provided with improved instructions, participants increased (and primarily employed) model-based learning during the two-stage task. We replicate some of these results. While we did not find evidence of pure model-based behaviour, improved instructions clearly promoted model-based behaviour compared to classical studies using other instructions. Therefore, our results suggest that participants deploy model-based strategies when they can form proper mental models of the two-stage task.

Contrary to da Silva and Hare (2020), our data revealed a significant main effect of Reward. Although this has often been linked to model-free behaviour, da Silva and Hare (2020) pointed out that it could also be driven by a poor understanding of the structure of the task. In support for his conclusion, da Silva and Hare (2020) (see their spaceships task and computational modelling work) found evidence that the adoption of an incorrect model could give rise to a significant effect of Reward. However, we think that this does not provide a satisfactory explanation for the main effect of Reward that we found in our experiments. First, in our replication of the magic carpet task, the vast majority of our participants had a good understanding of the task (i.e. 1 participant out of n=24 [0.04%] was removed from the magic carpet study by da Silva & Hare (2020), and 6 participants out of 59 [1%] were removed from our Replica study). Second, we also included a different experimental group fixing the locations of response options, in contrast to the classical two-stage task, where response options swap positions randomly across trials (e.g. Daw et al., 2011; and also da Silva & Hare, 2020). The logistic regression analyses revealed a larger effect of Reward in this group than in the otherwise identical replica condition. This effect of Reward is difficult to explain as the result of an incorrect internal model of the task, as the instructions in the Replica and Fixed-locations conditions were the same, allowing for the same mental model of the task. Rather, our results suggest that fixing option locations probably triggered habitual processes, facilitating the association between specific stimuli and specific motor commands. In agreement with our results, using a different task, but manipulating the consistency of response mappings, Molinero et al. (2021) found that reward-related cognitive prioritization was stronger when a constant response pattern was kept. Additionally, Neal, Wood, Wu & Kurlander (2011) provided evidence that previously formed habits disappeared when the response pattern was changed.

Thus, our results suggest that the main effect of Reward, often taken as a marker for model-free behaviour in the two-stage task, seems to be related to S-R learning. This result is important because it has been questioned whether the two-step task really taps into the habitual component of behaviour. For instance, if so, then this marker should increase with training, and not the contrary—and that was the result found in the only study that has manipulated the amount of training using the two-stage task to date (Economides et al., 2015)—. Moreover, there is empirical evidence showing that such model-free parameter does not correlate with habit strength measured by the canonical outcome devaluation test (Gillan et al., 2015). To this evidence, we should add the da Silva & Hare (2020) study itself. All these results apparently conflict with our suggestion that we were measuring, to some extent, habit strength using the two-stage task. We would argue that the conflict can be explained. As we show in our study, to tap into the functioning of the habit system it is essential that the motor response remains the same from trial to trial, given the same discriminative stimulus. This aspect of the design was not present in da Silva & Hare’s (2020) study. Also, the null result from Gillan et al. (2015) could be produced because their habit task was insensitive to the functioning of the habit system; indeed, other learning tasks previously used for studying habits have shown a lack of sensitivity for detecting them when they were further tested (Buabang et al., 2022; de Wit et al., 2018; de Houwer et al., 2018). Economides et al. (2015) found that extended training promoted model-based behaviour. Because they did not use the improved instructions as in da Silva & Hare (2020), it seems reasonable that a number of participants started with a wrong model of the task. Thus, extended training allowed these participants to learn and apply the correct model; hence, for these participants, model-based behaviour could be only available at the end of training. Learning the correct model of the task through training might overshadow the effect of S-R learning on participants’ behaviour. Future research should investigate the effect of the amount of S-R learning on model-free parameters in a task with improved instructions—ideally by manipulating both factors’ instructions (classic vs improved) and the amount of learning (little vs extended).

Our results concern to habitual responses thought as specific motor patterns that are activated after an S. It is important to note that that is not the only conceptualization of the “R” in S-R habits. For instance, Gadner and colleagues understand these responses as “impulses to act” whereas the act itself can change from instance to instance (e.g., Gardner et al., 2015). Our conception is different and concerns low level specific motor commands to achieve a certain goal (e.g., Du et al., 2022; Yang et al., 2022).

Although more research is obviously needed, our results are probably the first to show evidence compatible with the hypothesis that the two-step task can be used to infer the activity of the S-R habit system. We should be cautious and analyze if the increased Reward effect in the Fixed-locations study could be produced by other mechanisms. In this line, it has been suggested that “habits” are goal-directed responses produced when the participants activate a wrong goal (the “goal replacement hypothesis”, Kruglanski & Szumowska, 2020; de Houwer et al., 2018; Moors et al., 2019; Moors & de Houwer, 2017)—that is, not really habits. Indeed, there is an active controversy regarding this theory, with data and theoretical points in favour (e.g. Lally et al., 2010; Kaushal & Rhodes, 2015) and against (e.g. Hardwick et al., 2019; Luque et al., 2019) it. So, could our increased Reward effect in the Fixed-locations study be produced by differential activation of wrong goals? That seems unlikely. This hypothesis works better in experiments in which the reward is devalued (Dickinson, 1985; Dickinson et al., 1995), or Stimulus-Response-Outcome (S-R-O) mappings suddenly change after stable reinforcement learning (Harwick et al., 2019). In those scenarios, a participant could erroneously think that the value of the O, or the S-R-O map, did not change, and produce a goal-directed response with the appearance of a habit. But there is not a clear way (that we are aware of) to apply the “goal replacement hypothesis” to the two-step task.

## 4. Conclusions

To sum up, the present study converges with recent studies (i.e. da Silva & Hare, 2020; da Silva et al., 2023) in that it provides evidence of increased model-based behaviour when participants are provided with improved instructions in the two-stage task. In addition, we found that the effect of Reward can be promoted through invariably linking specific stimuli to certain motor commands. This provides evidence that a two-stage task behavioural marker, usually linked to model-free learning, is linked to S-R learning—a result that has been elusive until now.

## 5. Materials and Methods

### 5.1. Pre-registration

The methods and analysis plan employed in this study were pre-registered. The pre-registered protocol is publicly available at https://osf.io/x9sya

### 5.2. Participants

Following the recommendations by Brysbaert (2019) about minimum sample size in psychological research, we set our minimum sample one-hundred participants to ensure properly inter-group comparisons in our experiments. In total, one-hundred-and-eighteen participants from the Autonomous University of Madrid (UAM) were randomly assigned to the Replica condition (9 males, mean age: 22.29 years ± 5.54 SD; 50 females, mean age: 20.04 years ± 1.53 SD) or to the Fixed-Locations condition (9 males, mean age: 19.35 years ± 0.86 SD; 50 females, mean age: 20.35 years ± 2.13 SD). The best 3 participants in each group obtaining the largest scores in the task received 25€. Procedures were approved by the UAM ethics committee, and participants signed an informed consent before taking part in the experiment and were treated in accordance with the Helsinki declaration. All of them had normal vision or vision corrected to normality.

### 5.3. Apparatus

Participants were tested in individual cubicles, each with a standard PC and a monitor. Stimuli were presented using MATLAB with Psychtoolbox extensions (Brainard, 1997; Pelli, 1997; Kleiner, Brainard, & Pelli, 2007). Responses were collected using custom keyboards.

### 5.4. Task

Despite a few, but significant differences, a similar two-stage task was employed in the Replica and Fixed-Locations studies. The experimental task replicated that of da Silva & Hare (2020), which in turn was similar to Daw et al. (2011), except for some minor changes. As in da Silva and Hare (2020), the task was supported by a cover story causally explaining each transition and nuisance, so that a good understanding of the structure of the task was ensured (see the “Magic carpet task description” section in da Silva & Hare, 2020). Participants played the role of musicians living in a fantasy land, and obtained gold coins by playing the flute for an audience of genies living inside magic lamps in two different mountains, the Blue and the Pink mountain. Each mountain held two genies and participants were told that each lamp, with each genie inside, had a symbol written (Tibetan character) with the genie’s name in the local language. When participants arrived at a given mountain they had to choose a lamp, pick it and rub it. If the genie inside was in the mood for music then he would come out, listen to a song and give a gold coin to the musician. On such occasions, a genie with a coin was displayed on top of the lamp just chosen for 1,5 sec. Otherwise, a crossed “0” was displayed also for 1,5 sec. Participants were told, however, that the genies’ interest in music might change over time. Furthermore, in the Replica study, they were told that the lamps in each the Blue and Pink mountains could swap their positions between visits to a mountain. This was because every time they picked a lamp, they might leave it in a different place. In the Fixed-Locations study, participants were not told anything in this regard. This is because lamps’ positions were fixed across trials (positions were counterbalanced across participants).

To go to a certain mountain, participants had to choose between two magic carpets which would bring them there. The carpets had previously been enchanted by a magician so that each would fly to a different mountain. They had symbols written on them in the local language that meant “Blue Mountain” or “Pink Mountain”, depending on the destination of each magic carpet. Normally, carpets flew to their destination (common transitions). However, on rare occasions (rare transitions), travelling to the mountain of destination was too dangerous due to strong winds happening there. On such rare occasions, magic carpets were forced to land in the remaining mountain. Once more, in the Replica study, carpets could change sides between trials due to musicians putting them down and unrolling them on a different side of the room. On the contrary, in the Fixed-Locations study carpets remained in the same position across trials (positions were counterbalanced across participants). No specification on the location of the carpets was given during instructions.

In short, the task had a general main structure consisting of two stages happening in each trial (See Figure 3). In the first stage, participants needed to choose between two magic carpets that took them to either of two mountains (second stage). 70% of the times, a given carpet would take the participant to its assigned destination (common transition). However, the remaining 30% of the times, the carpet would bring the participant to its non-assigned mountain of destination (rare transition). Which carpet most probably flew to which mountain was randomized across participants. The position of the carpets (left or right) in the first stage changed randomly across trials in the Replica study and remained fixed in the Fixed-Locations study. In the second stage, participants were presented with two more options or states (lamps) and needed to choose one. These also changed their position (left or right) randomly across trials in the Replica study and remained fixed in the Fixed-Locations study. Finally, each state had a reward probability that varied between trials through a Gaussian random walk (mean 0, SD .025; with reflecting boundaries at 0.25 and 0.75) so that ongoing learning was encouraged. A pool of 20 Gaussian random walks was generated out of which, for each subject, 4 different random walks were selected at random to represent the reward probabilities of the total 4 second-stage state options of the 2 possible mountains in the second stage.

**Figure 3.**
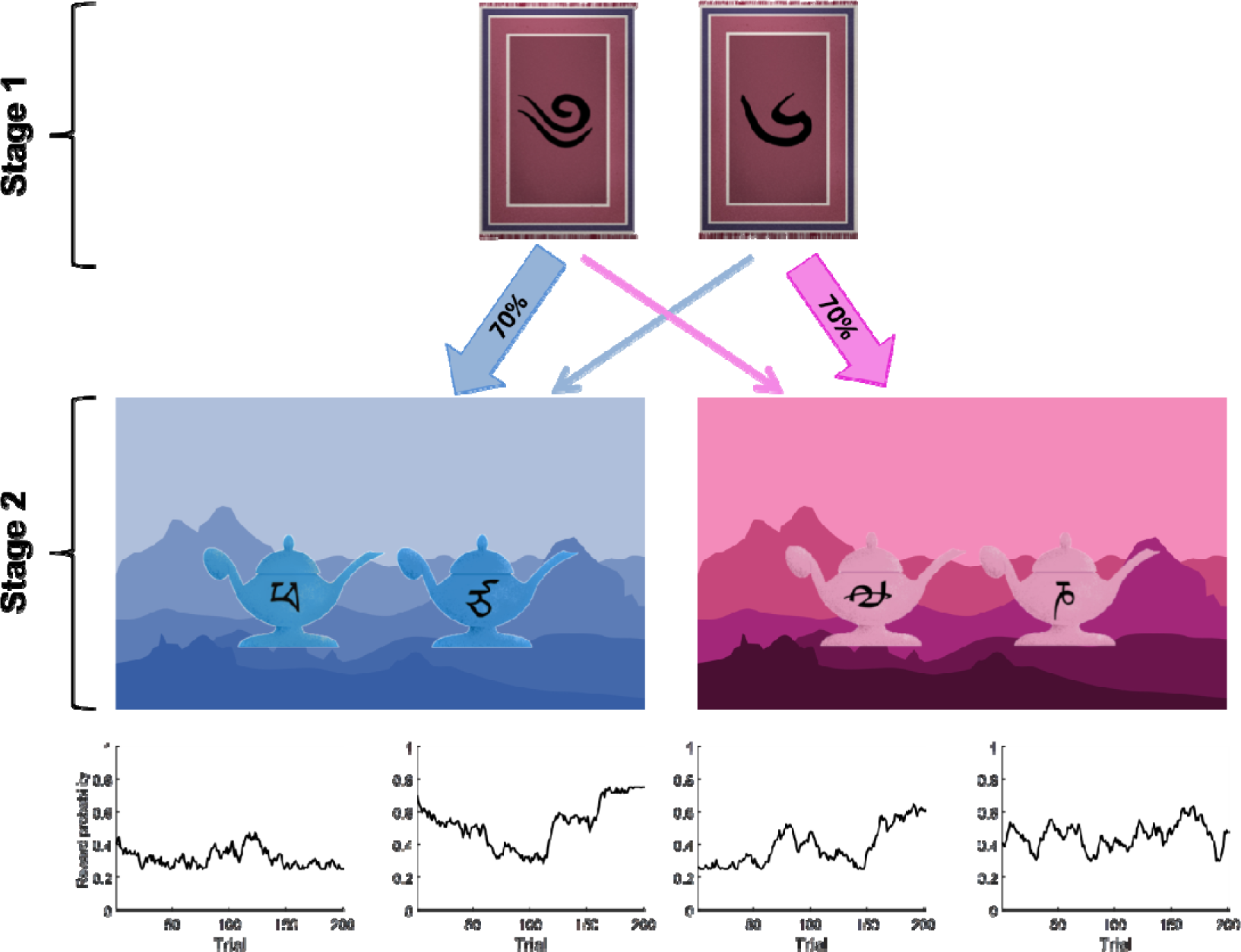
Two-stage task structure. During the first-stage state, participants need to choose between two options (magic carpets) that will bring them to two possible different states (Blue mountain [left] and Pink mountain [right]) in stage two. Transitions to second-stage states are probabilistic. A given magic carpet will transition to a given second-stage state with 70% probability, and it will transition to the remaining one with 30% probability. Once in a second-stage state, participants need to choose between two options (lamps). Each option has a reward probability that changes throughout trials by means of a Gaussian random walk (lower panels). Pictures of magic carpets and lamps are taken from da Silva & Hare (2020), and used in the studies of the present manuscript.

Participants were asked to always use the same finger for each response (left or right). That is, options on the left were selected using the left index finger, and options on the right were selected with the right index finger.

### 5.5. Procedure

The task consisted of 201 trials which were run along three blocks of 67 trials each. Participants were allowed to take a break in between blocks.

Choices in each stage were recorded, as well as response times (RTs). Participants were told that magic carpets in the first stage had to be chosen in less than 2 sec or else they would fly without them. In either case, lamps at the second stage had to be rubbed within 2 sec or the genies inside them would fall asleep and not come out. Trials in which participants failed to enter a response within 2 sec were be aborted with a message displayed on the screen: “TOO LATE! The magic carpets have flown without you” (for 7,5 sec) or “TOO LATE! The genies have fallen asleep” (for 1,5 sec). The duration of each message was such that the time spent was similar to the one that would have been spent if trials had not been aborted. We randomly selected the inter-trial interval from a uniform distribution ranging from 0.7 to 1.3 sec. During such interval, a Gaussian random noise mask (mean 0, SD 0.5) was presented to prevent possible visual aftereffects.

Previous to performing the task, participants completed 50 tutorial random flights. This was intended to make them aware of which transitions were common and which rare, and, in general, they became familiar with the narrative of the game and how to proceed in it. The only difference between the Replica and the Fixed-Locations group was that in the former participants were told that items may change places across trials (with an explanation of why that may happen) and in the latter they were not. During tutorial flights, participants were presented for 1 sec with a transition screen that made explicit to which mountain a carpet was flying. In addition, that screen showed them whether a carpet was flying to its mountain of destination (common transition) or was being flown away from it (rare transition). However, during the non-tutorial trials and because magic carpets were self-driving, musicians took a nap aboard it and only woke up upon arrival. A black screen was displayed during this period for 1 sec (not explicitly showing the mountain of destination and whether a transition had been common or not). Therefore, participants had to figure out the meaning of the symbols in each carpet for themselves. It is important to note that the mountains to which magic carpets flew during tutorial flights were different from the ones to which they flew during the main task. Namely, during tutorial flights, and in order not to interfere with the forthcoming task, magic carpets flew to the Black and Red mountains instead of the Blue and Pink mountains, where magic carpets flew during the main task. The magic carpets flying to each mountain and the lamps at the second stage used different Tibetan symbols from the ones used during the main task. A different pool of 20 Gaussian random walks for the second-stage state options’ reward probabilities was used for tutorial flights.

As explained above, the positions of the first and second-stage state options were randomized across trials in the Replica group and kept constant in the Fixed-Locations group but randomized across participants. The most likely transition through which each carpet flew to each mountain during tutorial flights was also randomized across participants.

Figure 4 shows the appearance and timing of the two-stage task both during tutorial flights (Figure 4A) and during the main task (Figure 4B).

**Figure 4.**
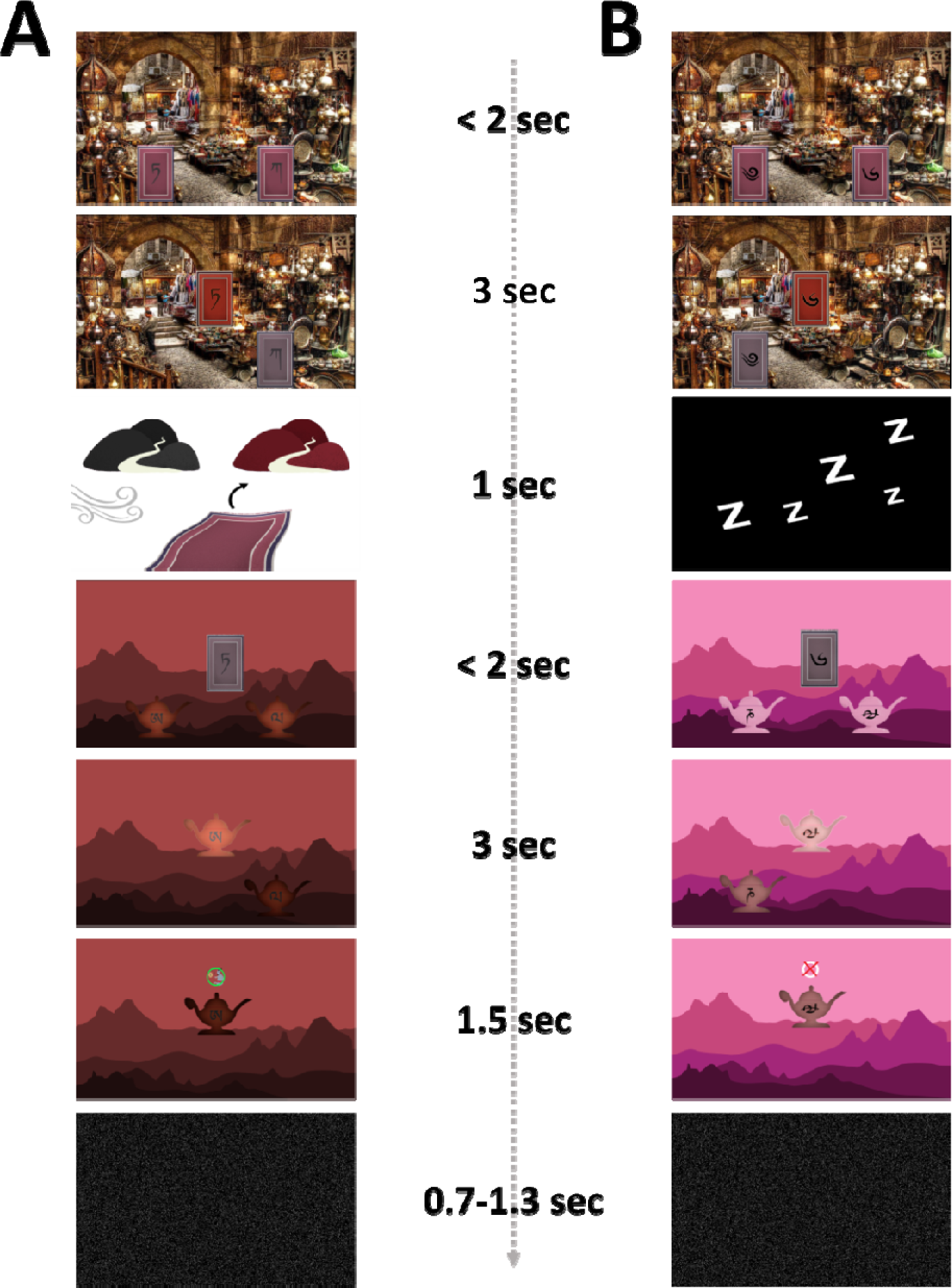
Appearance of the two-stage task and timing of events for tutorial flights (**A**) and the main task (**B**). First, one of two carpets needs to be selected in less than 2 sec, then the choice made is displayed for 3 sec. Afterwards, the transition to a given mountain is made (1 sec). In the case of **A**, such transition is shown to be a rare one. In **B**, this is not explicitly shown as the musician is taking a nap during the flight. Afterwards, once at a given mountain, a lamp needs to be chosen out of two different ones in less than 2 sec. Then the choice is displayed for 3 sec. Afterwards, a reward may be given (**A**) or not (**B**) depending on the interest in music at that moment of the genie inside the chosen lamp. The reward/non-reward stimulus is displayed for 1.5 sec. Finally, and right before the next trial, a Gaussian random noise mask is shown for 0.7-1.3 sec to prevent any visual aftereffects. (Note that during the main task musicians fly to the Blue and Pink mountains. However, during tutorial flights musicians fly to the Black and Red mountains) Pictures of magic carpets, lamps, genies and transition screens to a mountain are taken from da Silva & Hare (2020), and used in the studies of the present manuscript.

Importantly, before the tutorial flights, participants carefully read detailed instructions about them and completed a questionnaire with specific aspects about the task (See Appendix E). Wrong answers received feedback with the correct answer, making sure that participants did not start the tutorial flights without having understood the task. Also, after the 201 trials in the main task, participants were asked the following questions:

1) For each carpet symbol: What was the meaning of the symbol?

2) How difficult was the game? a) very easy, b) easy, c) average, d) difficult e) very difficult. (See Appendix D to check the number of participants reporting each difficulty category).

### 5.6. Analyses

First, in both experimental conditions (Replica and Fixed-Locations), trials in which participants had failed to enter a response within 2 sec were omitted (Replica: mean: 3.47 trials ± 5.87 SD; Fixed-Locations: 3.54 trials ± 4.26 SD). The pre-registration of the present study (https://osf.io/x9sya) specified that participants whose response time median absolute deviation (MAD) (Leys et al., 2013) was 3 points or larger would be excluded from analyses. The analyses reported in the main text do not remove participants based on this criterion. However, the reader can find analyses excluding them in Appendix B. The reason why we do not exclude these participants in the main manuscript is because this allows for a higher statistical power. Additionally, removing them does not significantly alter the results.

#### 5.6.1. Logistic regression analysis

A logistic regression analysis of consecutive trial pairs was performed separately for each participant in the Replica and Fixed-Locations studies. Trial pairs including a trial performed after a break during the task were excluded from analyses. The stay probability (i.e., the probability of repeating the same first-stage action as in the previous trial) was predicted as a function of two variables: reward (was the participant rewarded on the previous trial or not?) and transition (was the previous trial’s transition common or rare?) using the following equation:

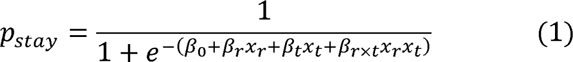

where *β*_0_, *β*_r_, *β*_t_ and *β*_r×t_ are, respectively, the coefficients for the intercept and the reward, transition and Reward × Transition effects. *x_r_* adopted values of +1 or −1 depending on whether the previous trial had been rewarded or not, and *x_t_* adopted values of +1 or −1 depending on whether the previous trial had had a common or a rare transition. The model was fit to each individual subject using the Matlab function “fitglm” in a way that coefficients were obtained for every participant.

A few participants in each experimental group (i.e., 6 in Replica and 6 in Fixed-Locations) provided responses in a very consistent manner. For instance, a subject may unequivocally choose the same first-stage state option when the previous trial had been rewarded and the transition had been common. In this scenario, perfect separation between classes occurs, making it impossible for Iteratively Reweighted Least Squares methods (as used by the Matlab function “fitglm”) to estimate parameter values. In these cases, we artificially changed at random only one of their choices producing a perfectly unequivocal pattern. With such consistent pattern broken, parameter estimation was made possible. We preferred this to remove participants where perfect separation of classes took place, as their cases were still informative about performance in the two-stage task. After all, this task may encourage such consistent patterns of behaviour.

#### 5.6.2. Hybrid reinforcement learning model fits

The standard hybrid reinforcement learning proposed by Daw et al. (2011), combining model-based learning with the model-free SARSA(λ) algorithm, was fitted to the empirical data of each individual participant taking part in the study.

At initiation (i.e., trial *t*= 1), the model-free (MF) values of the algorithm, 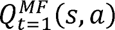 are set to zero. That is, 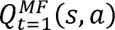 for each possible action *a* that an agent can perform in each stage, *s*, is 0. Once an action is chosen at the end of trial *t*, the 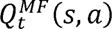 value for that action performed at a certain stage is updated. In the particular case of second-stage actions, *a*_2_, performed in a second-stage state, *s*_2_, (i.e. Pink and Blue mountains in Figure 3), 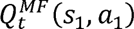, is updated through the following formula:

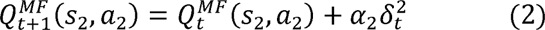

were *a*_2_stands for the learning rate for the second stage (0< *a*_2_ > 1) and 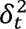 is the reward prediction error; namely, the current value of the action chosen, 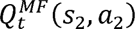, and the reward received, *r_t_* (0 or 1), and is defined as follows:

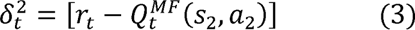

Regarding the chosen first-stage action, *a*_1_, at the first stage state *s*_1_, the value of the chosen action, 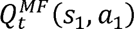, is updated as follows:

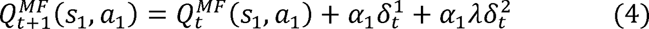

where 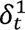 is the reward prediction error for the first stage, and is defined as:

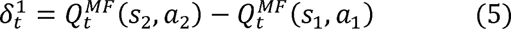

*a*_1_ is the learning rate for the second stage (0< *a*_2_ > 1) and λ is the eligibility parameter (0< λ> 1). This last parameter weights the effect of second-stage reward prediction error on first-stage action values.

Having explained the model-free (MF) values of the algorithm, we may now explain its model-based (MB) values. 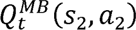 for action *a*_2_ at second-stage state *s*_2_ has the same meaning as the corresponding model-free value: 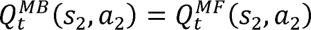. On the other hand, for each first-stage action, model-based values are calculated as follows:

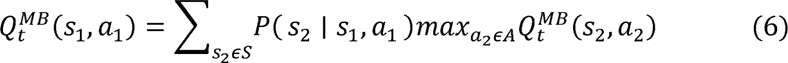

That is, model-based values for first-stage actions are computed when a decision is made from the values of second-stage actions, where *P*(*s*_2_ I *s*_1_, *a*_1_) stands for transition probability to state *s*_2_ through first-stage action, *s*_1_. *s* = {*pink,blue*} designates the possible second-stage states, and *A* designates the possible actions at those states.

Agents perform first-stage choices both based on model-free and model-based values according to a softmax distribution:

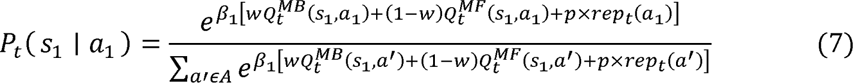

where *w* is a model-based weight whose value determines the amount of model-based influence (0

<*w*> 1). *β*_1_is the inverse temperature parameter for the first stage, and it models the exploration-exploitation trade-off during that stage. *p* is a perseveration parameter whose value has an effect on how prone agents are to repeating the previous trial’s first stage action in the next trial. Finally, *rep*_t_(*a*^r^) is a value defined as 1 if the first-stage action *a^’^*was performed in the previous trial (0 otherwise).

When it comes to the second stage, the probability of a given second-stage choice is computed as follows:

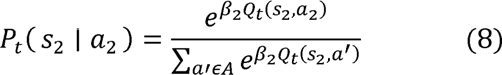

where model-free and model-based values for the corresponding second-stage actions are the same. This is because no tendency to repeat the previous action or keypress is assumed.

Estimates for model parameters, *a*_1_, *a*_2_, *β*_1_, *β*_2_, *w* and *p*, were obtained through maximum likelihood estimation. To this end, participants’ first-stage and second-stage responses as well as the transitions (common vs rare) that happened in each trial, together with the reward obtained were fed into the algorithm. In short, the reinforcement learning algorithm performed the same task as participants in a way that minimized the negative log-likelihood -log [*P_t_* (*s*_1_ *a*_1_)] to achieve each subject’s parameter values. For all participants, this process was repeated throughout 1000 iterations using varying starting values for the parameters. As in da Silva & Hare (2020), the model was coded in the Stan modelling language (Stan Development Team, 2012; Carpenter et al., 2017), and was further fitted to each subject’s data using the “cmdstanpy” library.

## 6. Acknowledgements

We would like to thank Carolina Feher da Silva for her help providing the code for reinforcement-learning model fitting analysis. As well, our version of the two-stage task took code from Github (https://github.com/DecisionNeurosciencePsychopathology/dom_conCog) and adapted it, for which we are grateful. Also, images of stimuli in the tasks were taken from those made publically available by da Silva and Hare (2020). RL was affiliated with the Department of Basic Psychology, Faculty of Psychology, Universidad Autónoma de Madrid, during the conceptualization and data collection of the study.

## Footnotes

1 When we designed our research, we expected participants in the Replica study to show mainly model-based behaviour, replicating this aspect of the results from da Silva & Hare (2020). This is acknowledged in the pre-registration protocol of this work (https://osf.io/x9sya). Therefore, we did not expect any differences between coefficients in any direction, and all the analyses in this regard are consequently two-tailed. However, when we compared the Replica study with the Fixed-Locations one, we did expect the Fixed-Locations study to detect a model-free component not present in the Replica study. Therefore, the t-test for the Reward coefficient is one-tailed. We also performed one-tailed tests on the Transition and Reward X Transition coefficients. This is because we expected a larger model-based component in the Replica study than in the Fixed-Locations one. The same logic was applied to all subsequent analyses.

## APPENDICES

### APPENDIX A

After participants had completed the two-stage task, they completed a questionnaire with key questions that can inform us about how well they understood the task. Here we report the results of the logistic regression analysis after excluding participants who failed any question from this questionnaire. In total, 6 participants were removed from the Replica condition (remaining sample size, n=53) and 6 participants were removed from the Fixed-Locations condition (remaining sample size, n=53). The same exclusion criterion was applied to the study developed by da Silva & Hare (2020), in which 1 participant did not answer correctly the final questionnaire. The remaining sample in this case was n=23. The results obtained did not differ significantly from the ones reported in the main manuscript using the full sample.

Figure A1 displays the results from the logistic regression analysis in a way similar to Figure 1 in the main manuscript.

**Figure A1.**
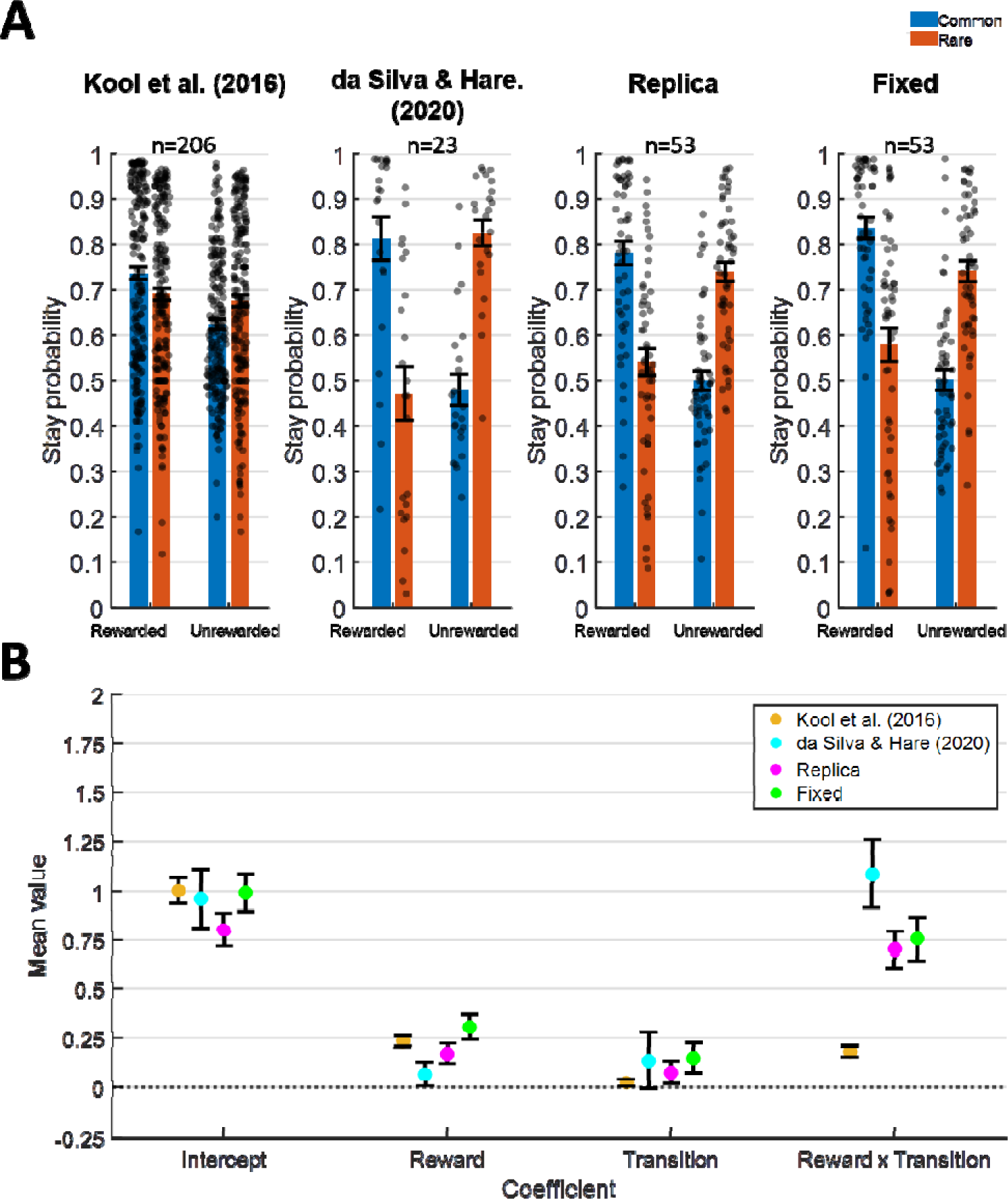
Results of the logistic regression analysis. **A.** Stay probabilities (probability of repeating the same response as in the previous trial) are shown in the cases when the transition in the previous trial had been Common (blue) or Rare (red). Results further distinguish whether the previous trial had been Rewarded or Unrewarded. The leftmost panel shows the results from the study by Kool et al. (2016) using the two-stage task paradigm by Daw et al. (2011); next, the results from the magic carpet experiment in da Silva & Hare (2020) are shown; the next panel shows the results for the Replica condition and the rightmost panel shows the results for the Fixed-Locations condition. Individual results are shown as well as the mean ± SEM. **B.** Coefficients for each of the logistic regression parameters, which were used to calculate the stay probabilities shown in the upper panels. The mean ± SEM is depicted.

Regarding comparisons between da Silva & Hare (2020) and our Replica study, statistically significant differences were observed in the Reward Transition coefficient, the one from da Silva & Hare (2020) being significantly larger (t(74)=-2.1161, p=0.0377, two-tailed, d=-0.5093, 95% CI [-0.7515, −0.0226]). No significant differences between the two were found in the rest of the parameters (Intercept: t(74)=-0.9596, p=0.3354, two-tailed, d=-0.2355, 95% CI [-0.4863, 0.1679]; Reward: t(74)=1.22427, p=0.2179, two-tailed, d=0.3203, 95% CI [-0.0627, 0.2706]; Transition: t(74)=-0.4632, p=0.6446, two-tailed, d=-0.1040, 95% CI [-0.3160, 0.1968]). Also, an analysis of which coefficient values statistically differ from zero showed a main effect of Reward x Transition but not of Reward in the study by da Silva & Hare (2020). In contrast, our Replica study showed a significant effect of both of the previous coefficients (See the one-sample t-tests in Table A1). A similar pattern can be appreciated in the study by Kool et al. (2016).

If we compare our coefficients from the Replica study with those by Kool et al. (2016), statistically significant differences were observed in the Reward x Transition coefficient (t(257)=7.1520, p<0.0001, two-tailed, d=0.9196, 95% CI [0.3744, 0.6590]). That is, model-based behaviour does experience an increase when improved instructions are provided. We did not find significant differences between our Replica condition and the study by Kool et al. (2016) in the remaining coefficients (Intercept: t(257)=-1.4609, p=0.1453, two-tailed, d=-0.2501, 95% CI [-0.4755,0.0705]; Reward: t(257)=-1.0791, p=0.2816, two-tailed, d=-0.1702, 95% CI [-0.1756, 0.0513]; Transition: t(257)=1.1677, p=0.2440, two-tailed, d=0.1550, 95% CI [-0.0370, 0.1450]). Our results comparing the Replica and Fixed-Locations conditions showed a significantly larger Reward parameter in the Fixed-Locations study (t(104)=-1.6612, p=0.0498, one-tailed, d=-0.3227, 95% CI (-∞, −0.0001]). No significant differences between conditions were found in the rest of the parameters (Intercept: t(104)=-1.4832, p=0.1410, two-tailed, d=-0.2881, 95% CI [-0.4477, 0.0646]; Transition: t(104)=-0.7868, p=0.7834, one-tailed, d=-0.1528, 95% CI [-0.2282, ∞); Reward x Transition: t(104)=-0.3757, p=0.6460, one-tailed, d=-0.0730, 95% CI [-0.2974, ∞)).

**Table A1.** Two-tailed one-sample t-tests for each of the coefficients, Intercept, Reward and Transition and Reward X Transition. Results are shown for all four studies, Kool et al. (2016), da Silva & Hare (2020), Replica and Fixed. The values of the t-statistics and their degrees of freedom are displayed, as well as p-values and 95% confidence intervals. p-values lower than 0.05 are marked in bold, indicating those cases where a coefficient significantly differs from zero.

|  |  | Kool et al. (2016) | dS & H (2020) | Replica | Fixed |
| --- | --- | --- | --- | --- | --- |
| Intercept | t(df) | t(205)=15.0920 | t(22)=6.4150 | t(52)=9.2916 | t(52)=10.3692 |
|  | p | **p<0.0001** | **p<0.0001** | **p<0.0001** | **p<0.0001** |
|  | 95% CI | [0.8744,1.1372] | [0.6513,1.2737] | [0.6298,0.9768] | [0.8024,1.1874] |
| Reward | t(df) | t(205)=8.9397 | t(22)=1.1381 | t(52)=3.6311 | t(52)=4.6660 |
|  | p | **p<0.0001** | 0.2673 | **0.0006** | **p<0.0001** |
|  | 95% CI | [0.1844,0.2887] | [-0.0580,0.1990] | [0.0780,0.2708] | [0.1772,0.4446] |
| Transition | t(df) | t(205)=1.3947 | t(22)=0.9599 | t(52)=1.3879 | t(52)=2.0755 |
|  | p | 0.1646 | 0.3475 | 0.1711 | **0.0429** |
|  | 95% CI | [-0.0105,0.0615] | [-0.1614,0.4395] | [-0.0354,0.1943] | [0.0051,0.3007] |
| Reward × Transition | t(df) | t(205)= 6.7859 | t(22)=6.3446 | t(52)= 7.4130 | t(52)=6.8141 |
|  | p | **p<0.0001** | **p<0.0001** | **p<0.0001** | **p<0.0001** |
|  | 95% CI | [0.1321,0.2402] | [0.7336,1.4462] | [0.5126,0.8931] | [0.5346,0.9809] |

### APPENDIX B

For each participant, we computed the mean response time, including responses in the first and second stages of the task, and the median absolute deviation (MAD) (Leys et al., 2013) of the average response times. Subjects whose MAD score was 3 points or larger were excluded from further analyses (i.e., logistic regression analysis and hybrid reinforcement learning model fits). In total, 2 participants were removed from the Replica condition (remaining sample size, n=57), 1 participant was removed from the magic carpet study by da Silva & Hare (2020) (remaining sample size, n=23), 6 participants were removed from the study by Kool et al. (2016), and no participants were removed from the Fixed-Locations condition. The results obtained do not differ significantly from the ones reported in the main manuscript using the full sample.

#### Logistic regression analysis

Figure A2 displays the results from the logistic regression analysis in a way similar to Figure 1 in the main manuscript.

**Figure A2.**
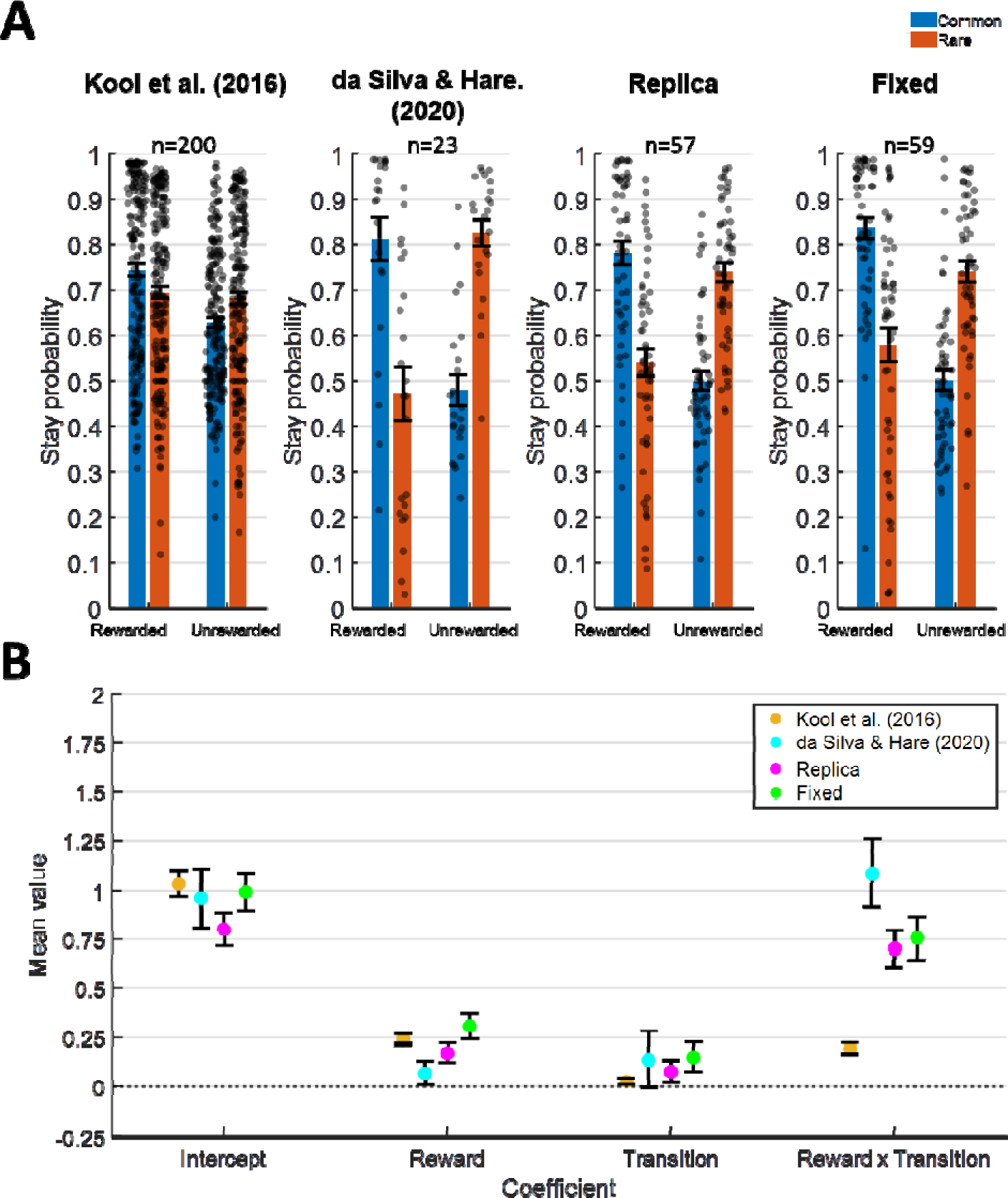
Results of the logistic regression analysis. **A.** Stay probabilities (probability of repeating the same response as in the previous trial) are shown in the cases when the transition in the previous trial had been Common (blue) or Rare (red). Results further distinguish whether the previous trial had been Rewarded or Unrewarded. The leftmost panel shows the results from the study by Kool et al. (2016) using the two-stage task paradigm by Daw et al. (2011); next, the results from the magic carpet experiment in da Silva & Hare (2020) are shown; the next panel shows the results for the Replica condition and the rightmost panel shows them for the Fixed-Locations condition. Individual results are shown as well as the mean ± SEM. **B.** Coefficients for each of the logistic regression parameters, which were used to calculate the stay probabilities shown in the upper panels. The mean ± SEM is depicted.

Regarding comparisons between da Silva & Hare (2020) and our Replica condition, statistically significant differences were found in the Reward Transition coefficient, the one from da Silva & Hare (2020) being larger (t(78)=-2.4427, p=0.0168, two-tailed, d=-0.5827, 95% CI [-0.8109, −0.0827]). No significant differences between the two were found in the rest of the parameters (Intercept: t(78)=-0.7665, p=0.4457, two-tailed, d=-0.1887, 95% CI [-0.4850, 0.2154]; Reward: t(78)=1.2978, p=0.1982, two-tailed, d=0.3290, 95% CI [-0.0557, 0.2644]; Transition: t(78)=-0.6192, p=0.5376, two-tailed, d=-0.1349, 95% CI [-0.3215, 0.1690]). Also, an analysis of which coefficient values statistically differ from zero showed a main effect of Reward x Transition but not of Reward in the study by da Silva & Hare (2020). In contrast, our Replica study showed a significant effect of both of the previous coefficients (See the one-sample t-tests in Table A2). A similar pattern was appreciated in the study by Kool et al. (2016).

If we compare our coefficients from the Replica study with those by Kool et al. (2016), statistically significant differences can be observed in the Reward x Transitioncoefficient (t(251)=6.9917, p<0.0001, two-tailed, d=0.9036, 95% CI [0.3643, 0.6499]). That is, model-based behaviour does experience an increase when improved instructions are provided. We did not find significant differences between our Replica condition and the study by Kool et al. (2016) in the rest of the coefficients (Intercept: t(251)=-1.6975, p=0.0908, two-tailed, d=-0.2907, 95% CI [-0.5071,0.0375]; Reward: t(251)=-1.1851, p=0.2371, two-tailed, d=-0.1877, 95% CI [-0.1830, 0.0455]; Transition: t(251)=1.1007, p=0.2721, two-tailed, d=0.1470, 95% CI [-0.0404, 0.1429]).

The comparison of the Replica and Fixed-Locations conditions reveals a significantly larger Reward parameter in the Fixed-Locations condition (t(114)=-1.8250, p=0.0353, one-tailed, d=-0.3399, 95% CI (-∞, −0.0127]). No significant differences between conditions were found in the rest of the parameters (Intercept: t(114)=-0.9993, p=0.3197, two-tailed, d=-0.1856, 95% CI [-0.3929, 0.1294]; Transition: t(114)=-0.9703, p=0.8330, one-tailed, d=-0.1896, 95% CI [-0.2239, ∞); Reward x Transition: t(114)=-0.4093, p=0.6584, one-tailed, d=-0.0761, 95% CI [-0.2898, ∞)).

**Table A2.** Two-tailed one-sample t-tests for each of the coefficients, Intercept, Reward and Transition and Reward Transition. Results are shown for all four studies, Kool et al. (2016), da Silva & Hare (2020), Replica and Fixed. The values of the t-statistics and their degrees of freedom are displayed, as well as p-values and 95% confidence intervals. p-values lower than 0.05 are marked in bold, indicating those cases where a coefficient significantly differs from zero.

|  |  | Kool et al. (2016) | dS & H (2020) | Replica | Fixed |
| --- | --- | --- | --- | --- | --- |
| Intercept | t(df) | t(199)=15.4403 | t(22)=6.4150 | t(56)=8.8127 | t(58)=10.3739 |
|  | p | **p<0.0001** | **p<0.0001** | **p<0.0001** | **p<0.0001** |
|  | 95% CI | [0.9051,1.1701] | [0.6513,1.2737] | [0.6395,1.0158] | [0.7743,1.1445] |
| Reward | t(df) | t(199)=9.0051 | t(22)=1.1381 | t(56)=3.9297 | t(58)=5.1248 |
|  | p | **p<0.0001** | 0.2673 | **0.0002** | **p<0.0001** |
|  | 95% CI | [0.1899,0.2964] | [-0.0580,0.1990] | [0.0857,0.2639] | [0.1912,0.4362] |
| Transition | t(df) | t(199)=1.5190 | t(22)=0.9599 | t(56)=1.1971 | t(58)=2.1822 |
|  | p | 0.1303 | 0.3475 | 0.2363 | **0.0332** |
|  | 95% CI | [-0.0084,0.0649] | [-0.1614,0.4395] | [-0.0423,0.1679] | [0.0120,0.2789] |
| Reward Transition | t(df) | t(199)= 7.0668 | t(22)=6.3446 | t(56)=6.8872 | t(58)=6.7266 |
|  | p | **p<0.0001** | **p<0.0001** | **p<0.0001** | **p<0.0001** |
|  | 95% CI | [0.1411,0.2504] | [0.7336,1.4462] | [0.4560,0.8302] | [0.4920,0.9089] |

#### Hybrid reinforcement learning model fits

Figure A3 displays the results from the Hybrid reinforcement learning model fits in a way similar to Figure 2 in the main manuscript.

**Figure A3.**
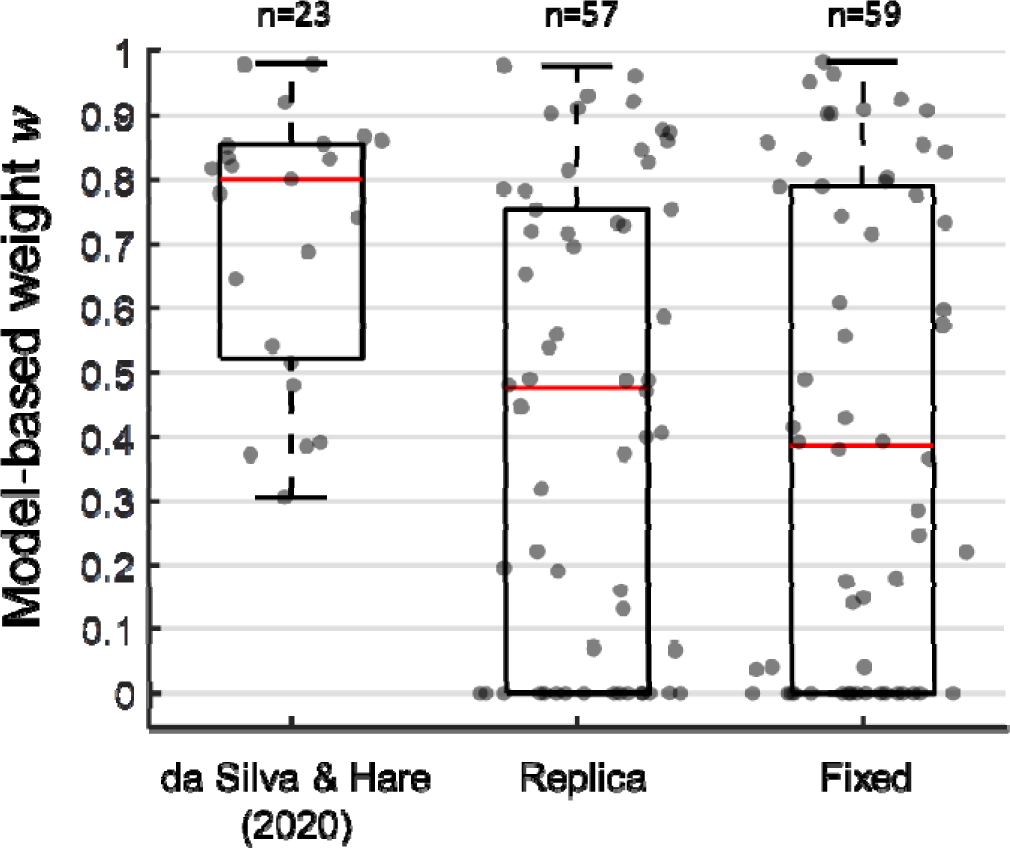
25%, 50% (median), and 75% percentiles of estimated parameters from the hybrid reinforcement learning algorithm as well as individual estimates. Data is shown for the *w* model-based weight parameter.

Our Replica condition fails to replicate the results from da Silva & Hare (2020) (: t(78)=3.5682, p=0.0006, two-tailed, d=0.9699, 95% CI [0.1243, 0.4380]). No statistically significant differences between the Replica and Fixed-Locations conditions exist (*w*: t(114)=0.1514, p=0.4400, one-tailed, d=0.0281, 95% CI [-0.1011,∞)).

### APPENDIX C

In Figure A4 we provide the estimates for all parameters in the standard hybrid reinforcement learning model proposed by Daw et al. (2011). We fitted this model to our Replica (n=59) and Fixed-locations (n=59) groups as well the data from the original magic carpet experiment by da Silva and Hare (2020) (n=24). Figure A4 also shows the negative log-likelihood, whose value was minimized during model fitting.

**Figure A4.**
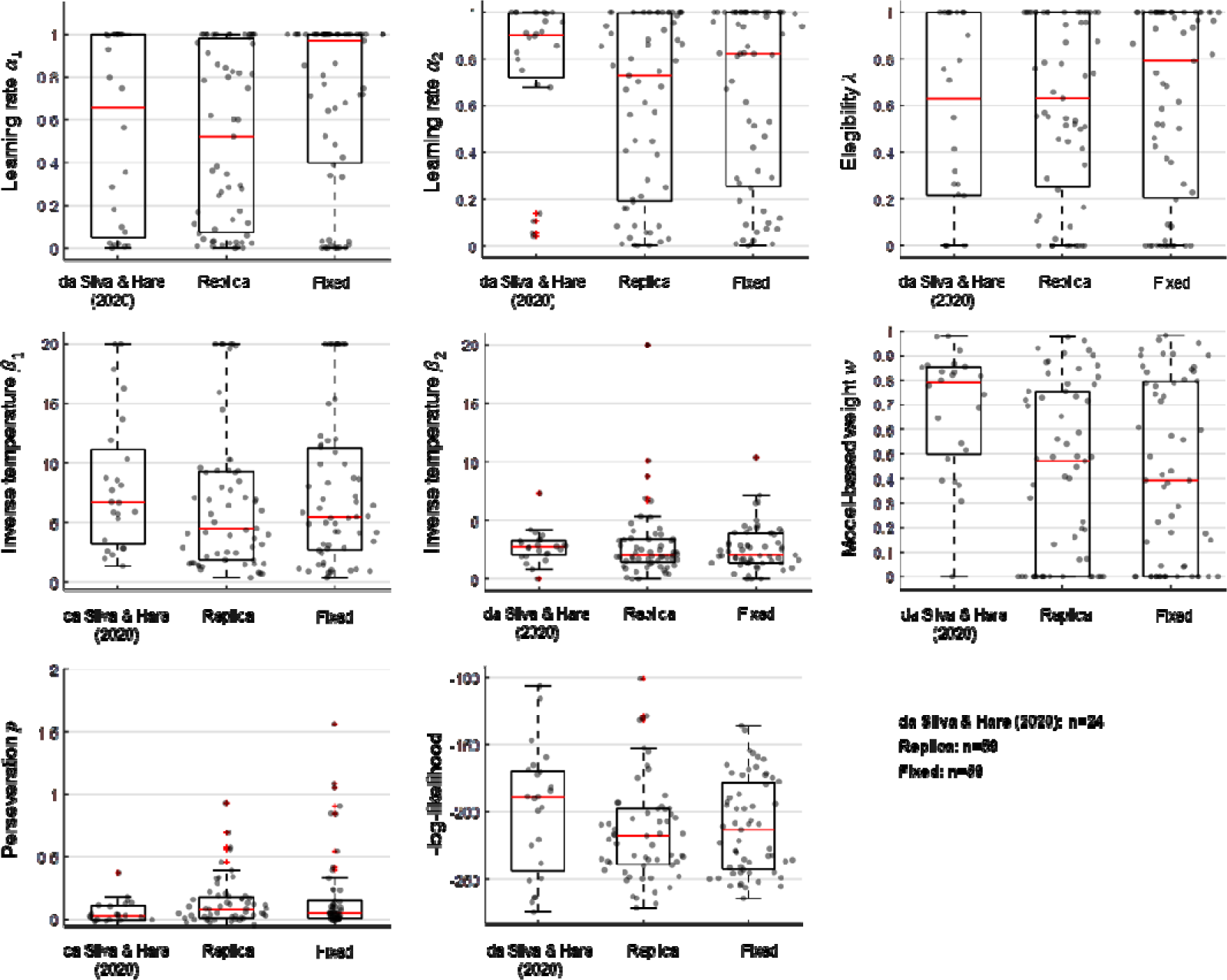
25%, 50% (median), and 75% percentiles as well as individual estimates of parameters and negative log-likelihood from the hybrid reinforcement learning algorithm. Red crosses indicate outliers. Data is shown for the and learning rate parameters, the elegibility parameter, the and inverse temperature parameters, the *w* model-based weight parameter, the *p* perseveration parameter and the negative log-likelihood.

### APPENDIX D

After the 201 trials of the main task, participants were asked about the difficulty of the task (“How difficult was the game? a) very easy, b) easy, c) average, d) difficult e) very difficult”). Figure A5 shows the number and proportion of participants reporting each difficulty category for the Replica (n=59) and Fixed-locations (n=59) groups as well as the magic carpet study by da Silva & Hare (2020).

**Figure A5.**
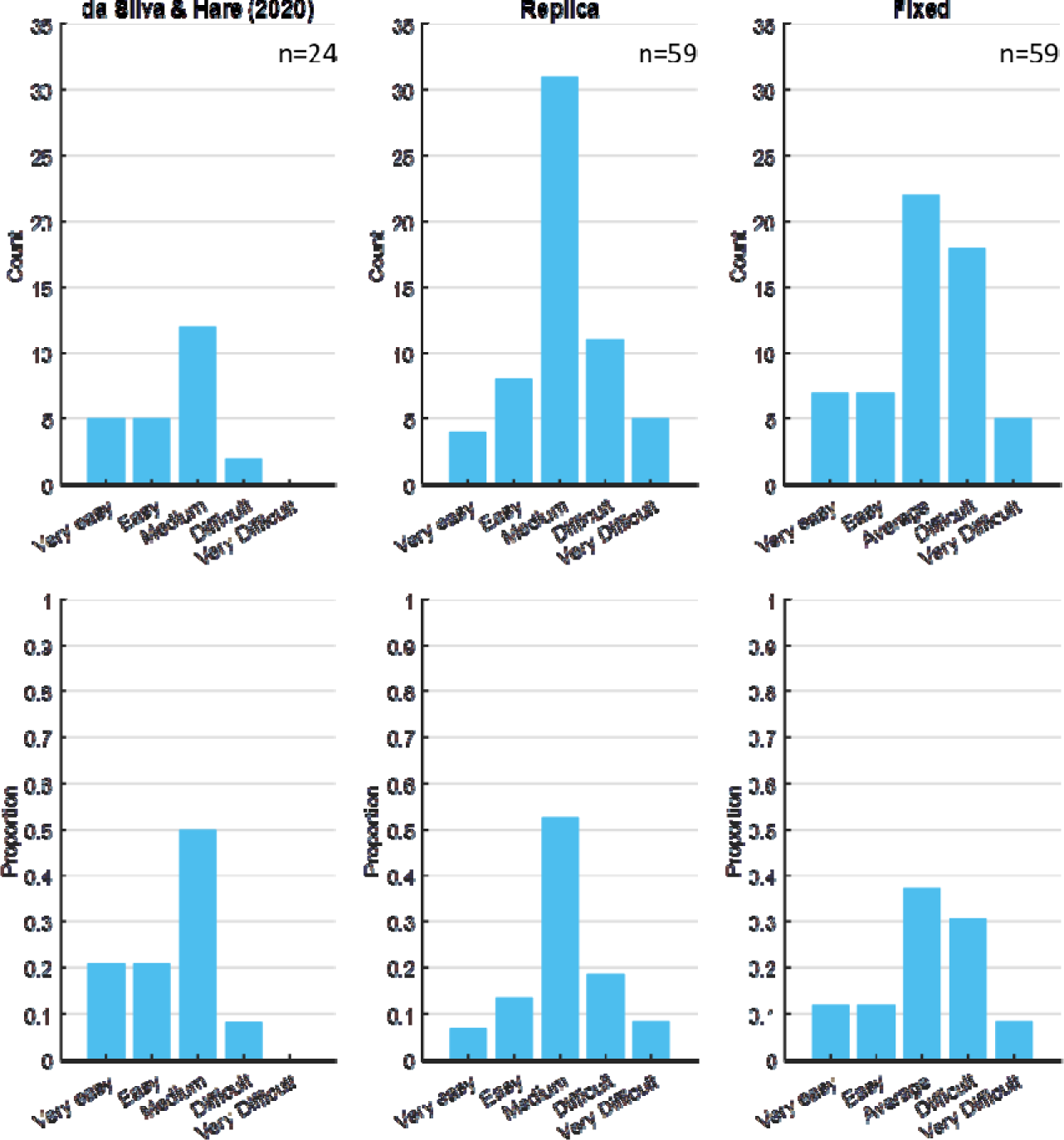
Number (upper panels) and proportion (lower panels) of participants in the magic carpet study by da Silva & Hare (2020), and the Replica and Fixed-locations groups reporting each difficulty category (“a) very easy, b) easy, c) average, d) difficult e) very difficult”) in the questionnaire after completing the 201 trials main task.

### APPENDIX E

Here we show the questionnaire that participants answered after reading the improved instructions on the two-stage task. Correct answers are marked in blue.

1. **Which mountain is most likely to have poor weather conditions?**

a. Both mountains may have bad weather conditions with equal probability.
b. The red mountain.
c. The black mountain.
2. **With which carpet is it most likely to have bad weather conditions?**

a. With the carpet enchanted to fly to the red mountain.
b. With the carpet enchanted to fly to the black mountain.
c. Both carpets may suffer bad weather conditions with equal probability.
3. **What does the symbol on the carpet you see mean? (e.g., to which mountain does that carpet fly when weather conditions are favourable?)** *(A magic carpet with a Tibetan symbol on it is shown on the screen)*.

a. Red mountain.
b. Black mountain. If a wrong answer is given, the correct answer is displayed: The symbol means “*mountain X*” and the carpet that carries it usually flies there. *(where “mountain X” is a given mountain, red or black)*.
4. **And what does the symbol of this other carpet mean? (e.g., to which mountain does that carpet fly when weather conditions are favourable?)** *(A magic carpet with the remaining Tibetan symbol not displayed in the previous question is shown on the screen)*.

a. Black mountain
b. Red mountain. If a wrong answer is given, the correct answer is displayed: The symbol means “*mountain Y*”, and the carpet that carries it usually flies there. *(where “mountain Y” is a given mountain, red or black)*.
5. **Where are you most likely to receive a coin from a genie?**

a. At the black mountain.
b. It depends on the interest in music of each genie.
c. At the red mountain.
6. **How can you tell if a genie is interested in music?**

a. When a genie is interested, it looks different.
b. Genies with a greater interest in music come out from their lamps more often to listen to music.
7. **How does the interest in music of a genie affect that of other genies?**

a. When a genie in a mountain has no interest, the other neither has it.
b. Each genie has its own interest in music, and it does not affect that of other genies.
c. When a genie in a mountain is interested, so is the other.
8. **How do bad weather conditions affect the interest in music of genies?**

a. Genies are more interested in music during windy days.
b. Genies live inside lamps, so they do not care about the weather.
c. Genies are more interested in music during windless days.

